# GWAS using 2b-RAD sequencing identified three mastitis important SNPs via two-stage association analysis in Chinese Holstein cows

**DOI:** 10.1101/434340

**Authors:** Fan Yang, Fanghui Chen, Lili Li, Li Yan, Tarig Badri, Chenglong Lv, Daolun Yu, Jie Chen, Chaofeng Xing, Jie Li, Genlin Wang, Honglin Li, Jun Li, Yafei Cai

**Author notes:** Co-first author: FAN Yang; Fanghui Chen. corresponding author: Yafei Cai; Jun Li. Other authors: Lili Li; Li Yan; Tarig Badrib; Chenglong Lv; Daolun; Jie Chen; Chaofeng Xing; Jie Li; Genlin Wang; Honglin Li.

## Abstract

**Background:** Bovine mastitis is a key disease restricting developing global dairy industry. Genomic wide association studies (GWAS) provided a convenient way to understand the biological basis of mastitis and better prevent or treat the disease. 2b-RADseq is a reduced-representation sequencing that offered a powerful method for genome-wide genetic marker development and genotyping. This study, GWAS using two-stage association analysis identified mastitis important genes’ single nucleotide polymorphisms (SNP) in Chinese Holstein cows.

**Results:** In the selected Chinese Holstein cows’ population, we identified 10,058 SNPs and predicted their allele frequencies. In stage I, 42 significant SNPs screened out in Chinese Holstein cows via Bayesian (P<0.001), while logistic regression model identified 51 SNPs (P<0.01). Twenty-seven significant SNPs appeared simultaneously in both analytical models, which of them only three significant SNPs (rs75762330, C>T, PIC=0.2999; rs88640083, A>G, PIC=0.1676; rs20438858, G>A, PIC=0.3366) located in non-coding region (introns and intergenic) screened out associated with inflammation or immune response. GO enrichment analysis showed that they annotated to three genes (PTK2B, SYK and TNFRSF21), respectively. Stage II? case-control study used to verify three important SNPs associated with dairy cows mastitis traits in independent population. Data suggested that the correlation between these three SNPs (rs75762330, P<0.025; rs88640083, P<0.005; rs20438858, P<0.001) and mastitis traits in dairy cows were consistent with stage I.

**Conclusion:** Two-stage association analysis approved that three significant SNPs associated with mastitis traits in Chinese Holstein cows. Gene function analysis indicated that three genes (PTK2B, SYK and TNFRSF21) involved in inflammation and immune response of dairy cows. Suggesting that they as new candidate genes have an impact on mastitis susceptibility (PTK2B and SYK, OR>1) or resistance (TNFRSF21, OR<1) in Chinese Holstein cows.

## Background

Bovine mastitis is the most complex and costly disease with high incidence, which seriously affects developing dairy industry worldwide (MAUNSELL *et al.* 1998; SCHUKKEN *et al.* 2009; WELDERUFAEL *et al.* 2017). Infection with mastitis causes direct economic losses in several ways, including dramatically discount in milk yield, treatment costs, condemnation of milk because of antibiotic or bacterial contamination. Also, higher than spontaneous elimination rates as well as, occasionally death of milk producer cows(SWINKELS *et al.* 2005; HALASA *et al.* 2007; HALASA *et al.* 2009; HOGEVEEN *et al.* 2011). Therefore, despite improvements in the breeding of disease-resistant cows, mastitis continues to be a notable challenge and the major profitable issue for dairy farmers. Previous studies reported that cow mastitis was a complex quantitative trait affected by multiple reasons, including genetic features, pathogen infections (HERTL *et al.* 2014; MOOSAVI *et al.* 2014; USMAN *et al.* 2015; POKORSKA *et al.* 2016; KIKU *et al.* 2017). It’s confirmed that bovine milk somatic cell count (SCC) or log-transformed SCC (somatic cell score, SCS) are the primary trait for detection of mastitis and have high hereditary capacity (WANG *et al.* 2015). Thus, Screening and identifying susceptibility or resistance genes associated with mastitis traits will improve the properties of dairy cow populations and is worthwhile to reduce the incidence of mastitis (SAHANA *et al.* 2014; KADRI *et al.* 2015; WANG *et al.* 2015). Different research strategies successfully used to identify significant genes associated with the mastitis traits, including SNP in a candidate gene, quantitative trait loci (QTL) and GWAS (BRONDUM *et al.* 2015; POKORSKA *et al.* 2016; ZHANG *et al.* 2016).

GWAS provides a convenient way to understand the biological basis of disease and better prevention or treatment (VISSCHER *et al.* 2017). In the past decade, it has been extensively in screen candidate gene mutagenesis to improve population productivity and disease resistance traits (DAETWYLER *et al.* 2014; CRISPIM *et al.* 2015; SAOWAPHAK *et al.* 2017; SELIMOVIC-HAMZA *et al.* 2017; VARSHNEY *et al.* 2017). It also widely regarded as a potential molecular marker assisted selection method based on SNPs in dairy cattle mastitis traits (WIGGANS *et al.* 2009; WANG *et al. 2015*). At chromosome level, GWAS data showed that Bos Taurus autosome 2, 4, 6, 10, 14, 18, and 20 associated with clinical mastitis significantly correlated with somatic cell scores in cows (SODELAND *et al.* 2011; MEREDITH *et al.* 2012; WIJGA *et al.* 2012). Besides, GWAS has arisen as one of the primary strategies in finding genetic variations associated with the traits. And many genetic associations have determined for a wide variety of prevalent, complex diseases as described in the GWAS list (HINDORFF *et al.* 2009). Sahana and his colleagues reported two clinical mastitis candidate genes (vitamin D-binding protein precursor, GC and neuropeptide FF receptor 2, NPFFR2) using high-density single nucleotide polymorphic array and WGAS (SAHANA *et al.* 2014). These two candidate genes detected to associate with mastitis traits in dairy cows through genomic sequencing in 2016 (ZHANG *et al.* 2016). In 2015, Wang et al. identified another two mastitis susceptibility genes (TRAPPC9 and ARHGAP39) in Chinese Holstein (WANG *et al.* 2015). However, Wu et al. detected five mastitis susceptibility genes (NPFFR2, SLC4A4, DCK, LIFR and EDN3) in Danish Holsteins (WU *et al.* 2015). Genetic variations in immune response, specific pathogen (LY75, DPP4, ITGB6 and NR4A2) and lymphocyte antigen-6 complex genes (LY6K, LY6D, LYNX1, LYPD2, SLURP1 and PSCA) might lead to clinical mastitis in American Holstein cows (TIEZZI *et al.* 2015). Additionally, single gene polymorphisms (CXCR1, MAP4K4) and their signaling pathways (TLR4/NF-κB) served as genetic markers for mastitis in different cow populations (POKORSKA *et al.* 2016; BHATTARAI *et al.* 2017). These results suggested that genetic variations or polymorphisms associated with mastitis traits are inconsistent, should screen and validated in different populations.

2b-RADseq, considered as a simplified and flexible restriction site-associated DNA (RAD) genotyping method based on IIB restriction endonuclease, provides a powerful method for identifying gene SNP in the population genome. It has strong technical repeatability, uniform depth of sequencing, high cost-effectiveness and genome coverage (WANG *et al.* 2012; GUO *et al.* 2014). Furthermore, the 2b-RADseq technique successfully predicted multilocus sequence typing (MLST) as well as provide more detailed on the population information than MLST technique. Therefore, the cost-effective and timesaving analysis strategy provided for large-scale studies on molecular epidemiology, public hygiene, systematic bacterial genetics, population genetics and bio-safety (PAULETTO *et al.* 2016; HERNANDEZ-CASTRO *et al.* 2017). Also, this method also suitable for erecting high-density genetic or linkage maps of genomic region or locus markers and revealing the regions associated with related traits by QTL mapping and association analysis (JIAO *et al.* 2014; ZHAO *et al.* 2017). More importantly, 2b-RAD can gain many SNPs through deep sequencing with fewer samples, and then identify the candidate genes related to traits (LUO *et al.* 2017). Therefore, 2b-RAD may be an ideal genotyping platform for screening mastitis resistance or susceptibility genes in dairy cattle.

In this study in order to identify mastitis susceptibility or resistance SNPs in Chinese Holstein and better understand the genetic and biological pathway of mastitis. We carried out: 1) 2b-RAD sequencing technique to sequence the whole genome for dairy cattle. 2) Identified and RADtyping SNPs. 3) GWAS analyzed the significant SNPs associated with mastitis traits via logistic regression analysis models. 4) Case-control study validated significant SNPs in independent dairy population. Then identified mastitis susceptibility or resistance SNPs, and evaluated the potential value of their associated genes in traits of Chinese Holstein cows.

## Methods

### Sample libraries and preparation

The experimental Chinese Holstein cows were from two different pastures of the same Dairy Company (Nanjing Weigang Dairy Co., Ltd.). Forty dairy cows selected from 596 lactating Chinese Holstein cows, which divided into two subgroups according to their clinical mastitis phenotypes: case group (20 cows) and control group (20 cows). 383 cows screened from 886 lactating cows in another pasture, with 73 in case group and 310 in control group. In their respective pastures, all animals have the same growth and feeding environment, similar production levels, the equivalent parity and lactation period.

Blood sampling performed using the tail vein blood sampling minimizes damage to cows (Firstly, the cows fixed in the column holders and the tails exposed outside the frame. Secondly, the blood collectors grasped the cow’s tails and lifted it upwards; they sterilized by alcohol cotton balls at the depressions at the midpoint of the 4th and 5th tail vertebrae. Then the tube blood collector penetrated the tail vein vertically to draw blood. Finally, after the blood drawn, the needle’s eye area pressed with a cotton ball for 30 seconds to fix the hemostasis and release the cows). Genomic DNA extracted from whole blood using *TIANamp Genomic DNA Kit*. The quality of genomic DNA detected by *NanoDrop* and *Agraros Gel* methods (extracted 3 microliters of genomic DNA, loaded on 1% agarose gel, 100 V CV 25 Minutes, viewed under ultraviolet light and photographed.).

### 2b-RAD library and sequencing

Forty sample libraries set up met a protocol developed by 2b-RAD sequencing needs with a little change and five-label tandem technique (RUBIN *et al.* 2010; WANG *et al.* 2012). The *Bos Taurus* genomes (ftp://ftp.ncbi.nlm.nih.gov/genomes/all/GCF/000/003/055/GCF_000003055.6_Bos_taurus_UMD_3.1.1/GCF_000003055.6_Bos_taurus_UMD_3.1.1_genomic.fna.gz) used as the reference for predicting electronic-enzyme-cut digestion of genomic DNA. Finally, *Bael* enzyme selected to digest genomic DNA. The restriction enzyme digested DNA fragment tags of each sample linked by standard 5‘-NNN-3’ connector. Paired-end sequencing carried out on the *Illumina Hiseq Xten* (https://support.illumina.com/downloads/sequencing-analysis-viewer-software-v2-18.html) platform after the quality control of the library was up to standard. Constructed the library followed WANG *et al.* (WANG *et al.* 2016a)(Figure S1) and the steps of the modeling included: (1) enzymatic digestion: ≥ 200ng genomic DNA digested by IIB restriction endonuclease; (2) Adding connectors: 5 different sets of connectors added to the digestion products respectively, with T4 DNA Ligase connection; (3) Amplification: T4 DNA Ligase connection products amplified by PCR; (4) Series: according to the information of 5 sets of connectors, the five labels connected in series; (5) Pooling: Barcode sequence added to the connection products and mixed library; (6) Sequencing: high-quality library that qualified, then on-machine sequencing.

### Raw Reads quality control

The original sequencing (Raw Data or Raw Reads) gained using the *Illumina HiSeq* sequencing platform. The *Phred value* was a role of sequence base error rate (Figure S2). It gained by calculating the probability model of prediction base recognition error base recognition. The calculation formula was: Q_*Phred*_ = −10log_10_(*phared*) (Table S1). Then deleted the reads contained the junction sequence and N base ratio ≥8% reads, got Clean Reads, then spliced by *Pear* (*Version 0.9.6*) software (http://pear.php.net/package/HTTP_WebDAV_Client/download/0.9.6/). Based on locating each sample, high-quality Enzyme Reads containing cleavage recognition sites extracted. *SOAP* (*version 2.21*) (http://soap.genomics.org.cn/soapaligner.html#down2) (Short Oligonucleotide Analysis Package) software used to align Enzyme Reads with the reference sequence (-r0 denotes unique comparison; -M4 represents optimal comparison; -v2 index comparisons allow two mismatched). Unique tags gained by the same Reads clustering showed the sequencing depth of the tags.

### SNPs genotyping and Linkage Disequilibrium (LD) analysis

SNP marker typing (RAD typing) performed on Enzyme Reads using the maximum likelihood (ML) method in SOAP software. The statistic SNP typing results, using R Package cluster analysis of the differences between sample SNPs. SNP annotated using SnpEff software (Version: 4.1g) (http://snpeff.sourceforge.net/) to determine located the SNPs in the gene and affected amino acid changes.

Plink software used to calculate the r^2^ value of the pairwise SNPs, with the main parameters set to: --r^2^-Id-window-kb 1000-Id-window 50-Id-window-r^2^ 0.2. According to the median of the software, the work F(x) = 1/ (log_10_ ((x+10^(7-C)^)/10^7^) + C) used to fit, and mapping by chromosome/grouping. To find the LD block in the case-control group, we add the parameters “block output GAB-pair wise Tagging’ to R package runner. Based on the 2b-RAD sequencing results, determined the optimal maximum distance for each LD block.

### Population structure and genetic diversity

After the 2b-RAD sequencing technology processed the samples, the PC analysis method used to evaluate the population. Then a correlation test performed for each subgroup, including the first five PC selected as covariate analysis signs for population division adjustment. The PC detailed data showed in Table S3. As far as we know, the first two influential feature vectors selected to draw the correlation between the samples. As shown in Figure S5 (a), case-control overlaps in two groups, no outliers detected, and it conformed to the rule of sample collection. To assess experimental samples’ genetic diversity, polymorphic information content (PIC), observed heterozygosity (Ho) and expected heterozygosity (He) values also calculated for each SNP site. In addition, the genetic differentiation coefficient between the subgroups was statistical.

### SNPs and mastitis traits association analysis

We considered a GWAS for the quality traits of dairy cow mastitis, and 2b-RAD markers genotype for each individual SNP locus. To ensure the accuracy of the analysis, we used multi-stage GWAS identified important SNPs associated with mastitis traits. Stage 1: we selected clinical mastitis and normal control Chinese Holstein cows for GWAS. We assumed that there is a true SNP tag associated with mastitis for each genotype in the genomic region. And calculated the correlation statistic for each SNP site and selected the strongest associated SNPs as the causal SNPs. If the region contains a single SNPs, then the most significant related SNPs is most likely a causal SNPs. Calculated the associated statistics of the SNP and identified significant SNPs. GWAS analysis performed using Bayesian and Logistic regression analysis model to compare important SNPs between case-controls. Quantile-Quantile Plot (QQ-plot) evaluated the rationality of the two analysis models. GO enrichment analysis performed on all genes with SNPs, and their functions described in conjunction with GO annotations. Hypergeometric Distribution Test (*Cytoscape* software) used to calculate significant gene enrichment in each GO entry. Stage 2: in another independent Chinese Holstein dairy population, validated these important SNPs screened in stage 1.

### Statistical model

#### Principal analysis method (PCA)

PCA is a method that uses for dimensionality reduction on data to study how to condense many original into a few factors with minimal information loss (LI *et al.* 2017). In this experiment, let F_1_ denote the main sub-index formed by the first linear combination of the original, F_1_=a_11_*X*_1_+ a_12_*X*_1_+…… +a_1m_*X*_m_, (m stands for the m*th* index). Information obtained by each principal component can measure by its variance. The larger the variance, the more information the F contains. If the first principal component is not enough to represent the initial m indicators, then consider selecting the second index F_2_. The existing data of F_1_ does not need to appear in F_2_ again. That is, F_2_ and F_1_ should be independent, irrelevant, and expressed by their covariance. And so on to build F_1_, F_2_… F_n_, as equation (1).

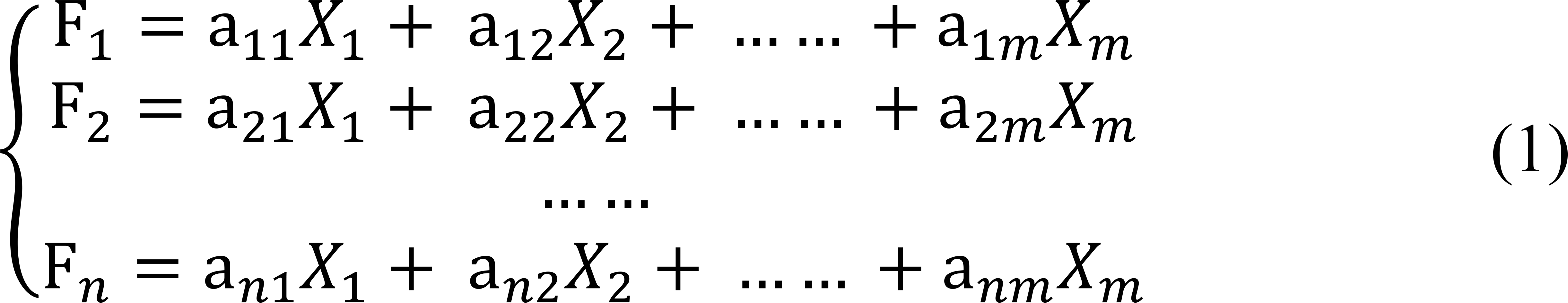

#### Bayesian and Logistic regression model association Analysis

Linear models are a common method for correlation analysis of phenotypes and genotypes. Strict quality control used to remove poorly performing SNP marker loci in RAD typing. Bayesian and Logistic regression model introduced for GWAS detected SNPs associated with clinical mastitis in dairy cows. First, built the following linear regression equation based on phenotype (GUO *et al.* 2018):

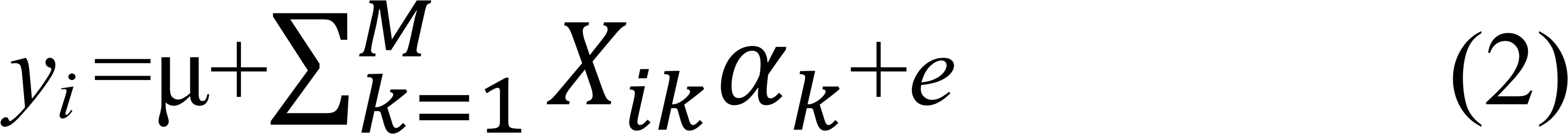

Where *y*_*i*_ is a vector of phenotype for individual *i*; M is the number of SNPs; μ is a vector of the overall mean of traits phenotypes; α_*k*_ is a vector of additive correlation effect of the *k*th SNPs; X*ik* is a vector of the genotype (0, 1, or 2) of the *k*th SNPs observed on the *i*th individual; and *e* is a vector of residual effect.

The Bayesian model assumed the SNPs effect was a prior normal distribution. Firstly, we consider the possibility that each SNP locus truly associated with the mastitis phenotype in GWAS. Select a value п for the prior probability H_1_. The correlation between SNPs and dairy mastitis traits quantified using п values. A pre-estimation of the SNPs truly associated with cow mastitis trait performed by a specific п value (10^−4^−10^−6^). While the probability of H_0_ considered to be (1-п). Secondly, calculated the Bayes factor for each SNPs. The Bayesian factor (BF) is the ratio between the probability of data at H_1_ and H_0_. Null assumption is H_0_ (*θ*_*het*_ = *θ*_*hom*_ = 0). H_1_, at least one *θ*_*het*_ =t_1_ and *θ*_*hom*_ =t_2_ value is non-zero (WELLCOME TRUST CASE CONTROL *et al.* 2012).

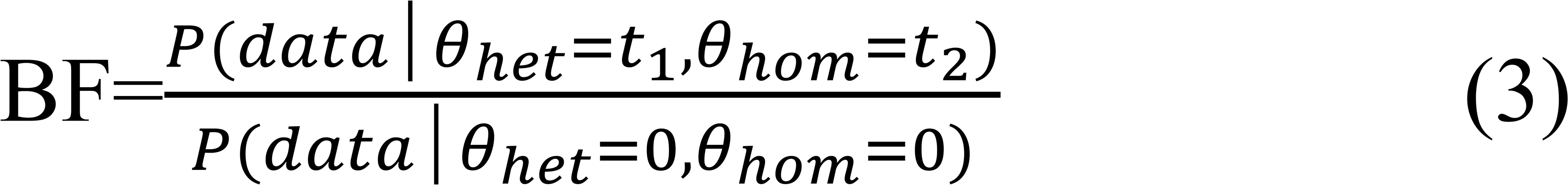

Where, *θ*_*het*_ is odds ratios (ORs) logarithm between the heterozygote and the common homozygote. *θ*_*hom*_ is the ORs logarithm between rare and common homozygotes. Then counted the posterior odds (PO) under the H_1_ condition: PO= BF×п/ (1-п). And posterior probability of association (PPA= PO/ (1+PO)) can be regarded as a Bayesian simulation of P value.

The SNPs effects variances were independent of each other, and each of which followed the same independent distribution (IID) as the inverse chi-square prior normal distribution where *v* is a parameter of the degree of freedom and *S*^*2*^ the parameter of scale:

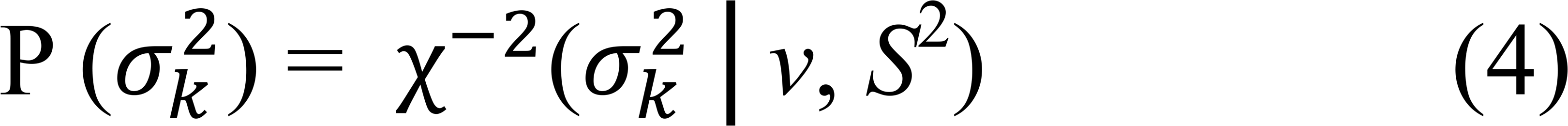

A prior distribution of the criticality of each SNP effect was a t-distribution (MEUWISSEN *et al.* 2001; GUO *et al.* 2018):

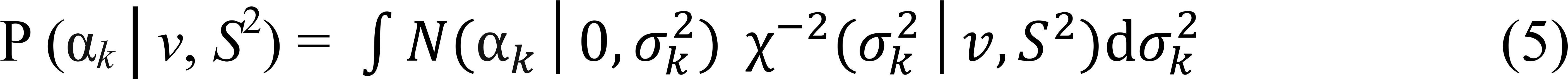

The prior for *a*_*k*_ depends on the variance of each SNPs, and each variance has an inverse Chi-square. SNP has null effect with probability п or is a normal distribution with probability (1-п), 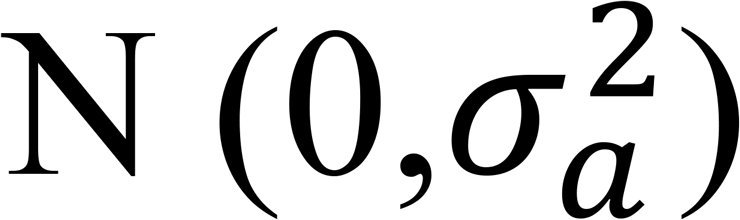 (GIANOLA 2013):

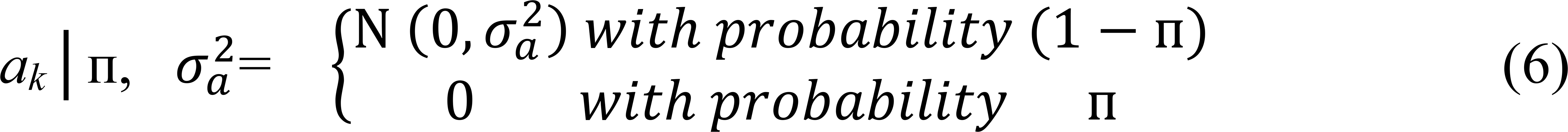

Where 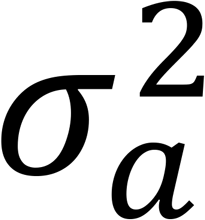 is represents the common variance of all non-zero SNPs effects, and it prorated prior distribution of the chis-square, 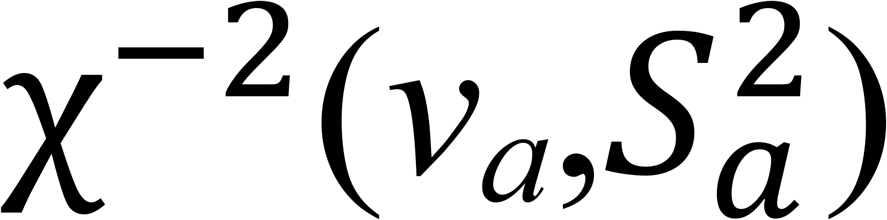. The unknown п value in the model predicted from its prior distribution (considered as uniform between 0 and 1) or п− uniform (0, 1).

*v*_*a*_ is designated as 4, 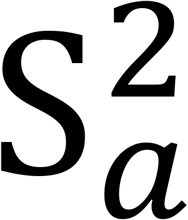 is calculated by additive variance.

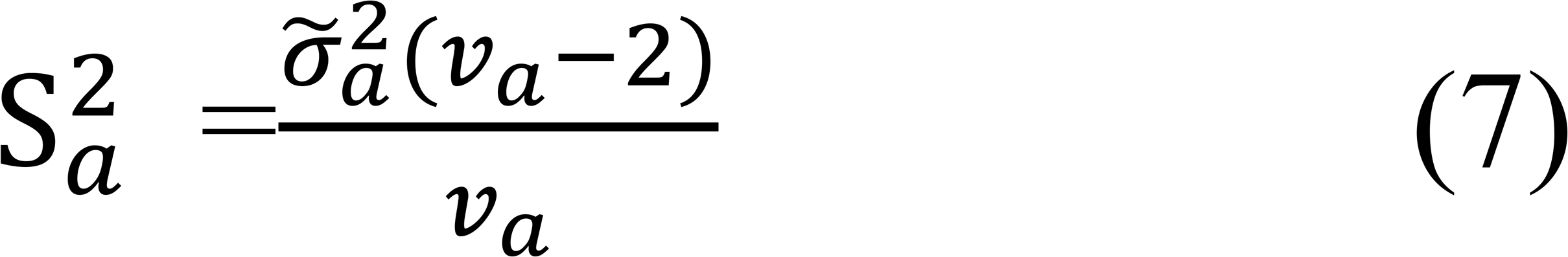

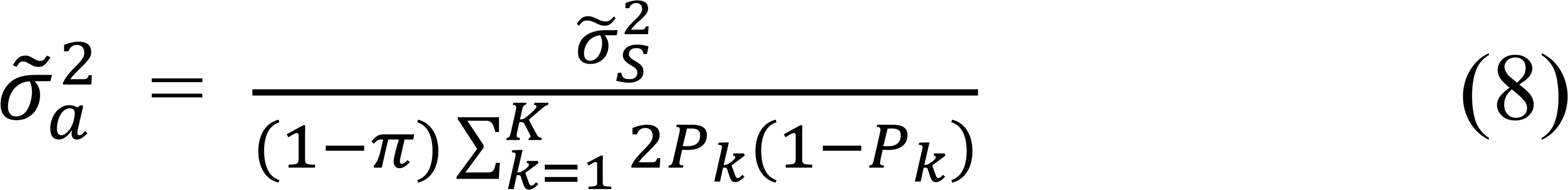

Where the allele frequency of the *k*th SNP is *P*_*k*_ the variance of a given tag is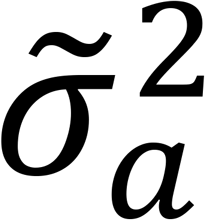, and the additive genetic variance 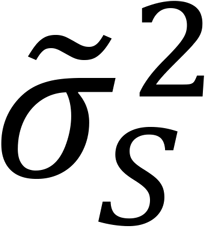 is elucidated by SNPs.

Then assuming that SNPs affects the mastitis phenotypic traits, we constructed logistic regression equation to predict SNPs associated with clinical mastitis in dairy cows. And we established a fitted logistic regression equation (BISCARINI *et al.* 2016; WANG *et al.* 2016b):

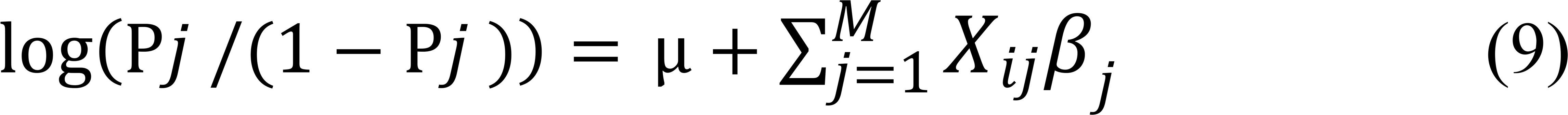

Where, P_*j*_ is the probability of occurrence of clinical phenotype under a condition X_*ij*_; (1-p_*j*_) is the probability that phenotype does not occur; X_*ij*_ = (X_*1j*_, X_*2j*_, X_*3j*_…… X_*mj*_) is the genotype of individual i at position j (0, 1, or 2); *β*_*j*_ is the effect of *j*th SNPs; and m is the number of samples; μ is the overall mean of traits phenotypes.

In the logistic regression model, Y= (μ+Σβ_i_X_i_) or (log *P*/(1 −*P*)), the equation can transform into another equation form:

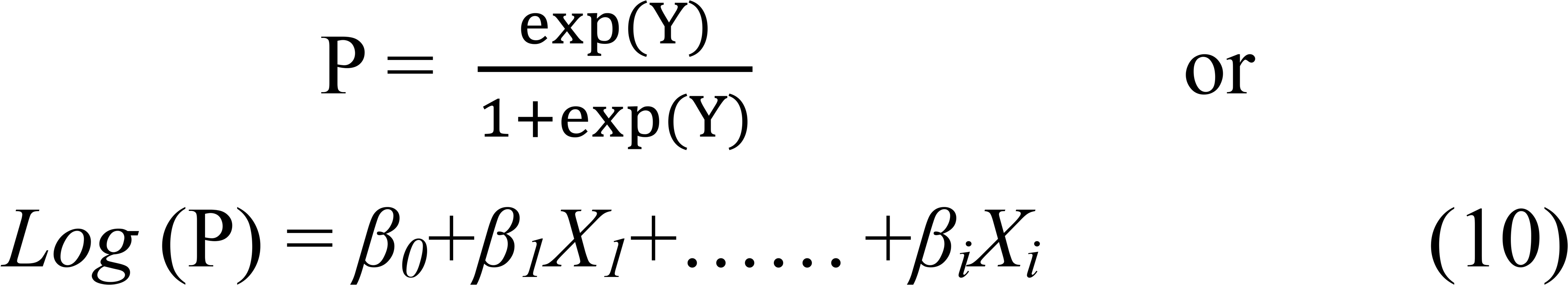

Where P is clinical mastitis phenotype, *X*_*i*_ is the genotype of individual *i*, *β*_*i*_ is the odds ratio (OR). The equation of expression between P and variable *X*_*i*_ can derive by equation transformation:

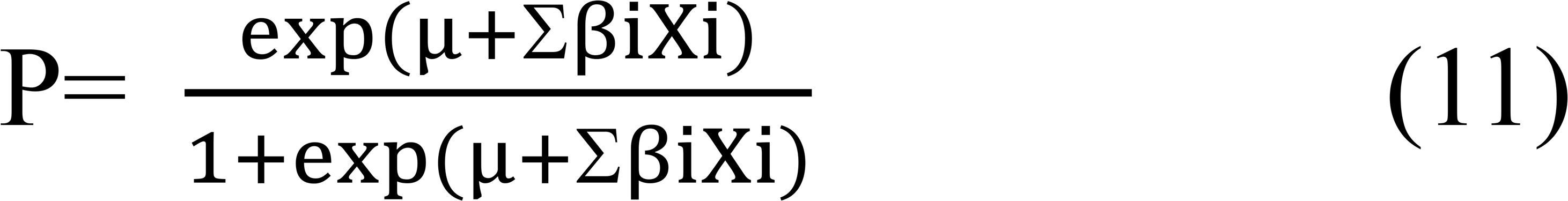

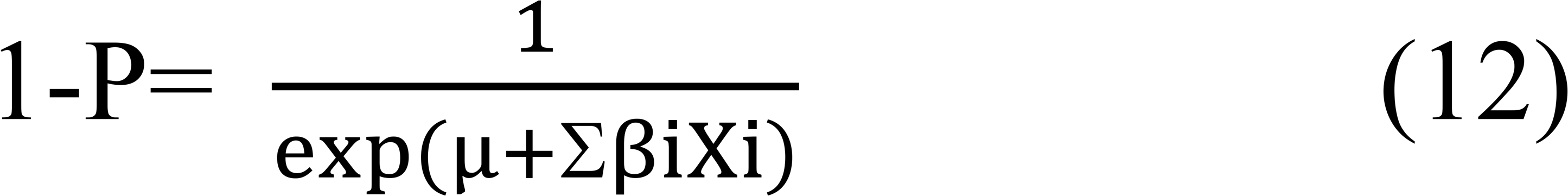

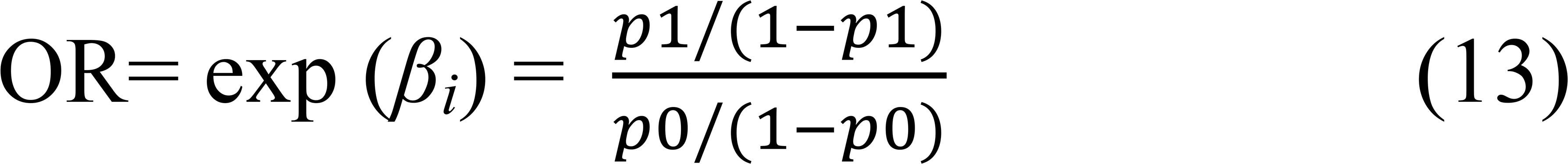

The greater the value of *β*_*i*_, the greater the influence of Y. 95% confidence interval: CI= exp (*β*_*i*_±1.96SE (*β*_*i*_)).

#### Case-control population verification analysis

We determined the number of validation samples using a matching design and case-control unequal (case/control=1/h).

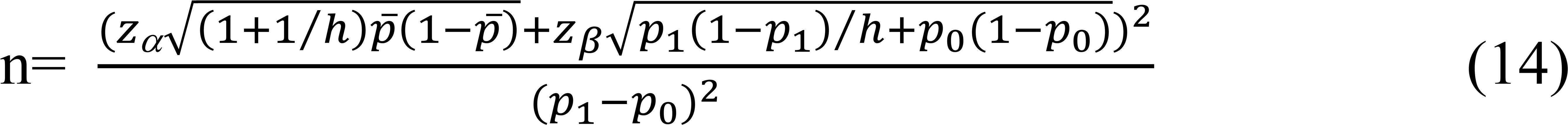

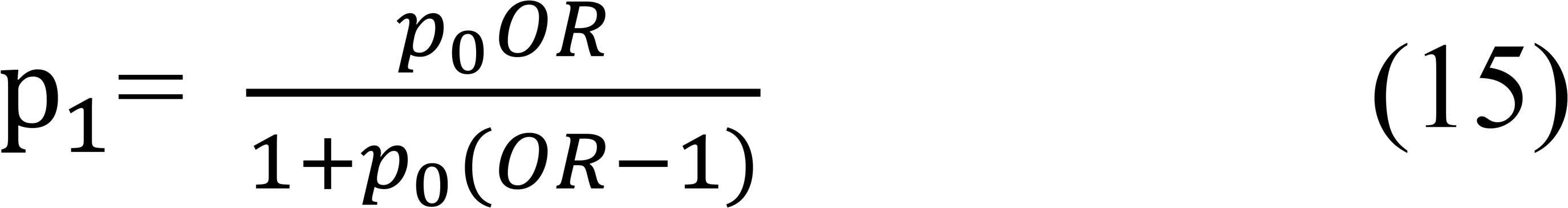

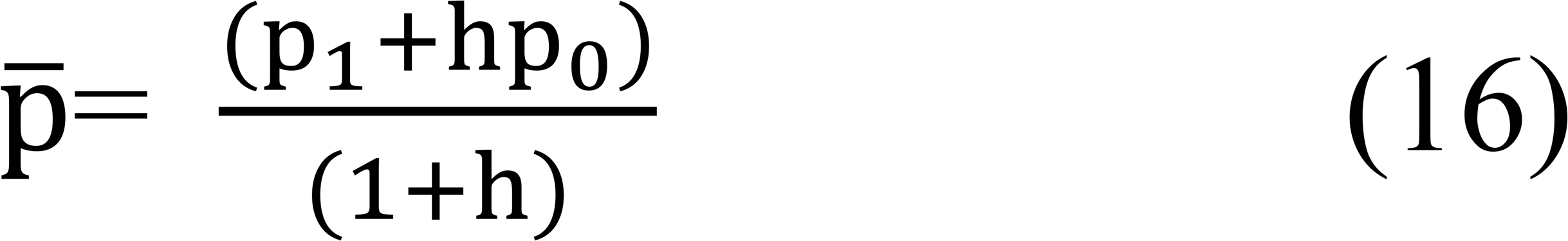

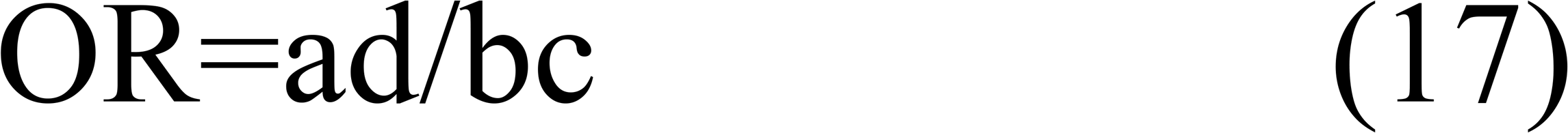

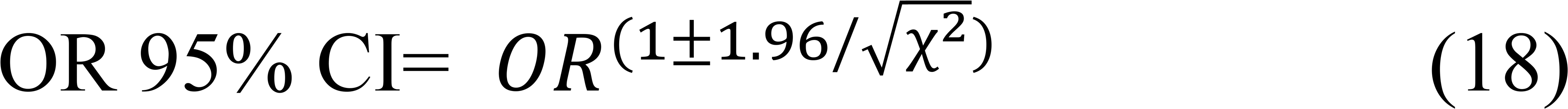

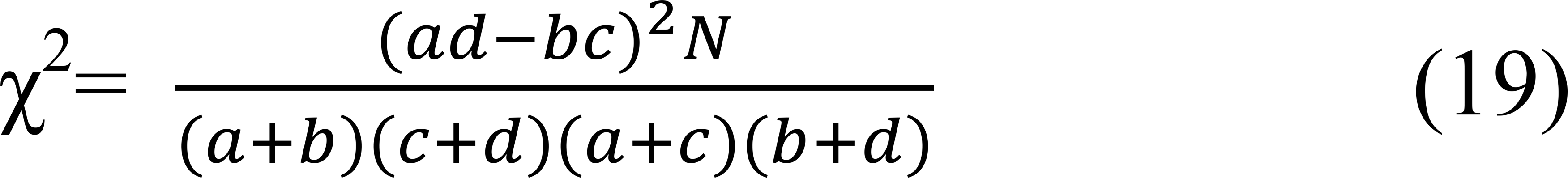

Where, n is the number of cows in clinical mastitis. *N* is the total number of cows in verification population. P_0_ is the exposure rate of SNPs in the control group. P_1_ is the exposure rate of SNPs in the case group. OR is the exposure ratio (Odds ratio). α is the probability of hypothesis testing type I errors. β is the probability of hypothesis type II errors and (1− β) is the expected test assurance. OR 95% CI is 95% confidence interval.

The Attributable Fraction reflects the probability that a case will be randomly selected from the population due to the SNPs.

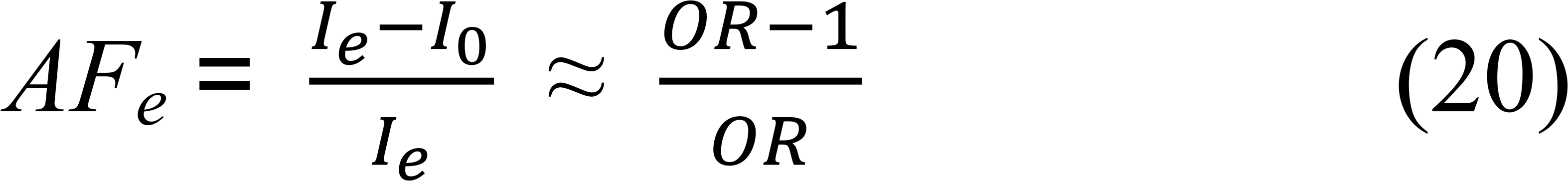

Where, *I*_*e*_ is the incidence of the site mutation group; *I*_*0*_ is incidence of the site non-mutation group. Incidence is generally not available in case-control studies and only OR obtained. *AF*_*e*_ refers to the proportion of mastitis caused by the SNPs to all mastitis.

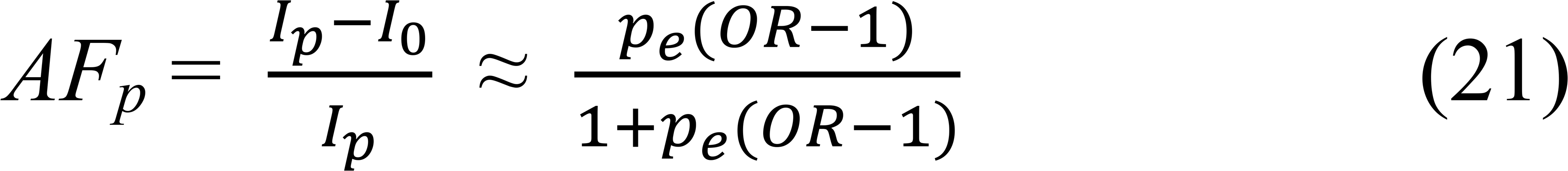

Where, AF_p_ indicates the proportion of mastitis caused by the SNPs in all mastitis. I_p_ is the total incidence of mastitis in Chinese Holstein cows. I_0_ is the incidence of mastitis with non-mutation at the SNP locus. P_e_ is the mutation rate of SNP locus in control group.

## Results

### Restriction endonuclease digestion and unique tags statistics

In this project, *Bael* restriction endonuclease used to digest genomic DNA of Chinese Holstein cows. Cluster analysis performed on the same read to gain a unique tag for each sample, and calculated the depth of sequencing for each tag. Removing the tag with a sequencing depth of less than 3×, the average number of tags each sample was 198,948 and the average sequencing depth was 17.43× (Figure 1, Third Ring). The average tag spacing between tags was about 9589 bp (Figure S3) and the unique tags alignment ratio for all samples was 59.69% ~ 72.71%.

**Figure 1.**
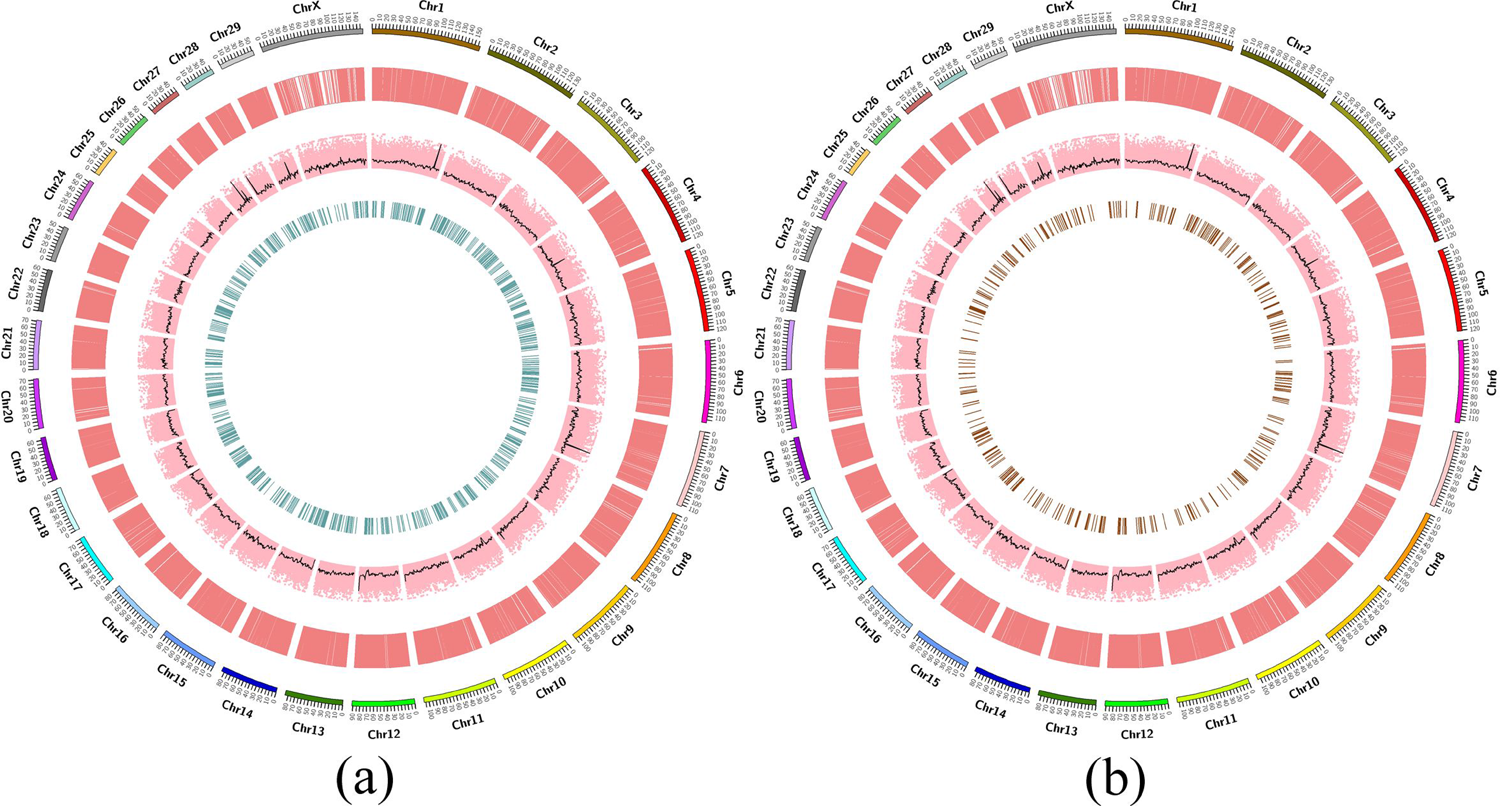
Schematic diagrams of all SNPs at chromosomal location cricos distribution and sequencing depth summary (a): Bayesian model analysis of SNPs quality traits in all samples (from outer to inner). The outmost circle was the chromosome scale; the second was the differential SNPs for all samples; the third was the average sequencing depth for each of the 40 samples at each locus (pink, if the depth over 50, count by 50) and the average depth of sequencing within 1 M window (black line, depth exceeds 50, count by 50); the fourth ring was the SNPs of the quality traits Chi square P< 0.05. (b): Logistic regression analysis of SNPs quality traits in all samples. The first to third rings were the same as (a). The fourth loop was the result of logistic regression analysis of differences in SNPs (P< 0.05).

### SNPs RADtyping and genotype

After RADtyping and filtering, 10,058 SNPs (Figure 1, Second Ring) screened out for all samples. The distribution of SNPs on the chromosomes of each sample was based on sliding window statistics (Figure S4). Then we counted all the SNP locus allele frequencies. Cochran-Armitage test analyze association between single SNPs genotypes and case-control status showed in Figure S5(b). Here, since heterozygote risk assessment intermediated between two homozygotes, this line fit the data reasonably which matched to additive genotype risk. In this case there was no deviation, and the test was convincing. And details gave in Table S2. At the same time, we counted the genetic differentiation coefficient (Fst= 0.01869) between two groups.

The range of Fst value between groups was 0-1. Fst value was close to 0, which indicated that the genetic differentiation between the two groups was smaller.

### Genome Wide Association Analysis

The SNPs associated statistic obeys the multivariate normal distribution. We also calculated the likelihood of possible causal states of the SNP. Each SNP has two potential causal: effect or no effect of the SNP. Therefore, for a possible subset of each, we need to consider the 2^n^ likelihood of the SNP. For each of these states, a multivariate normal distribution used to calculate the probability of the data for a given causal state. Thus, to identify the best SNPs set, a large amount of computations must be performed. The data showed that analysis of the two models was a slightly different in quality traits for all SNPs. Bayesian analysis screened 42 significant SNPs when P<0.001 (Table S4), while logistic regression analysis model identified 51 SNPs under P<0.01 condition (Table S5). Under the above P-value conditions, 27 significant SNP sites appeared simultaneously in the two analytical models (Table 1). As expected, significant SNPs screened in the two analytical models under their P-value conditions will vary, respectively. The QQ-plot (Quantile-Quantile Plot) evaluated the rationality of the two statistical models. Figure 2 showed that P value observations consistent with expected values at all SNP site, indicated that the two analysis models were reasonable. In the upper-right corner of Figure 2(a), candidate sites with high significance and potential associated with mastitis traits. However, in Figure 2(b), SNPs that significantly associated with mastitis traits were not apparent. This might be related to the fact that cow mastitis controlled by micro-multiple genes; gene effects too weak, or the sequencing population size we selected was insufficient.

**Figure 2.**
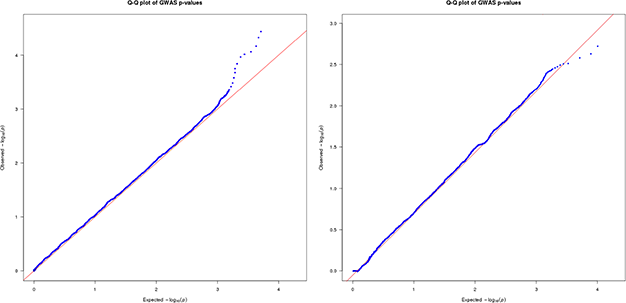
Q-Q plots (Quantile-Quantile plots) diagram for consistency of the observed and predicted values of SNPs P-value. (a) and (b) represented the consistency of Bayesian and logistic regression analysis for SNPs observed and expected value – log10 (P), respectively. (a) There were SNPs P values exceeded expected, which suggested that these locus might be significantly associated with dairy cows’ mastitis traits. (b) The P value observation is almost the same as the expected, indicated that the analysis model was reasonable.

In Bayesian analysis model (Table S4), The OR and 95% CI values of the SNPs rs21068792 were all “Na”, there was a missing value. The U95 of the 6 SNPs (rs98302192, rs49124945, rs57070376, rs13685463, rs57506421 and rs58979699) were “Nan”, which means meaningless number. The OR value of these 11 SNPs (rs114843903, rs38937721, rs5881560, rs17514753, rs17518215, rs22015301, rs77887746, rs9704351, rs20438858, rs26414259 and rs50888452) less than 1 indicated that these SNPs were protective reasons for the related phenotypes. The other 24 SNPs’ OR values greater than 1 indicated that these SNPs were risk for related phenotypes. With regard to logistic regression model, 51 SNPs marked when P<0.01(Table S5). A total of 22 SNPs’ OR value less than 1, and other 29 SNPs were greater than 1. We noticed that 8 (rs114843903, rs5881560, rs17514753, rs17518215, rs22015301, rs9704351, rs20438858 and rs5088452) SNPs’ OR values<1, while for Bayesian model as well. Table 1 also showed that 19 SNPs’ OR values in the two models were great than 1.

### SNPs GO annotations

We annotated all 27 significant SNPs to determine their location in the chromosomal genome. Table 2 showed that 14 SNPs located in the intergenic region, 10 in intron, and 1 in 3’-UTR, upstream and downstream, respectively. Except for the rs33866959 (A>T, transition) site, all other sites are transversion. The PIC value of rs86640083 less than 0.25 (low polymorphism), while the others all in 0.25 to 0.5 (moderate polymorphism). Go enrichment for 27 significant SNPs revealed that only 3 SNPs (rs75762330 (C>T, PIC= 0.2999), rs88640083 (A>G, PIC= 0.1676) and rs20438858 (G>A, PIC= 0.3366)) associated with immune role (Table 3, Figure 3). SNPs rs75762330 (C>T, OR>1, PIC= 0.2999> 0.25) in PTK2B gene located on BTA 8, and belonged to moderate polymorphism. PTK2B, also called Pyk2, regulates humeral and homeostatic cell homeostasis (RACIOPPI *et al.* 2012; KREMER *et al.* 2014; RHEE *et al.* 2014; LLEWELLYN *et al.* 2017). And rs88640083 (A>G, OR>1, PIC=0.1676<0.25) in intergenic nearby SYK gene located on BTA 8 and was low polymorphism. SYK is a non-receptor tyrosine kinase and considered as an important regulator factor for adaptive immunity and played a vital role in TLR4 signaling pathway (CHOI *et al.* 2015; SCHWEIGHOFFER *et al.* 2017). SNPs rs20438858 (G>A, OR<1, PIC= 0.3366>0.25) in TNFRSF21 located on BTA 23 and was moderate polymorphism. TNFRSF21, also known as Death receptor 6 (DR6), is a member of the TNF/TNFR family and played a critical role in immune response and inflammation (LOCKSLEY *et al.* 2001; STRILIC *et al.* 2016; FUJIKURA *et al.* 2017). Other 24 SNPs were statistically significant in both analytical models, but GO annotations showed that they did not have the function of inflammation or immune response Combining the two models provided support for comprehensive data from all SNPs, suggested that association between the 3 significant SNPs and the risk of mastitis in dairy cows based on conventionally accepted genome-wide statistical significance thresholds.

**Figure 3.**
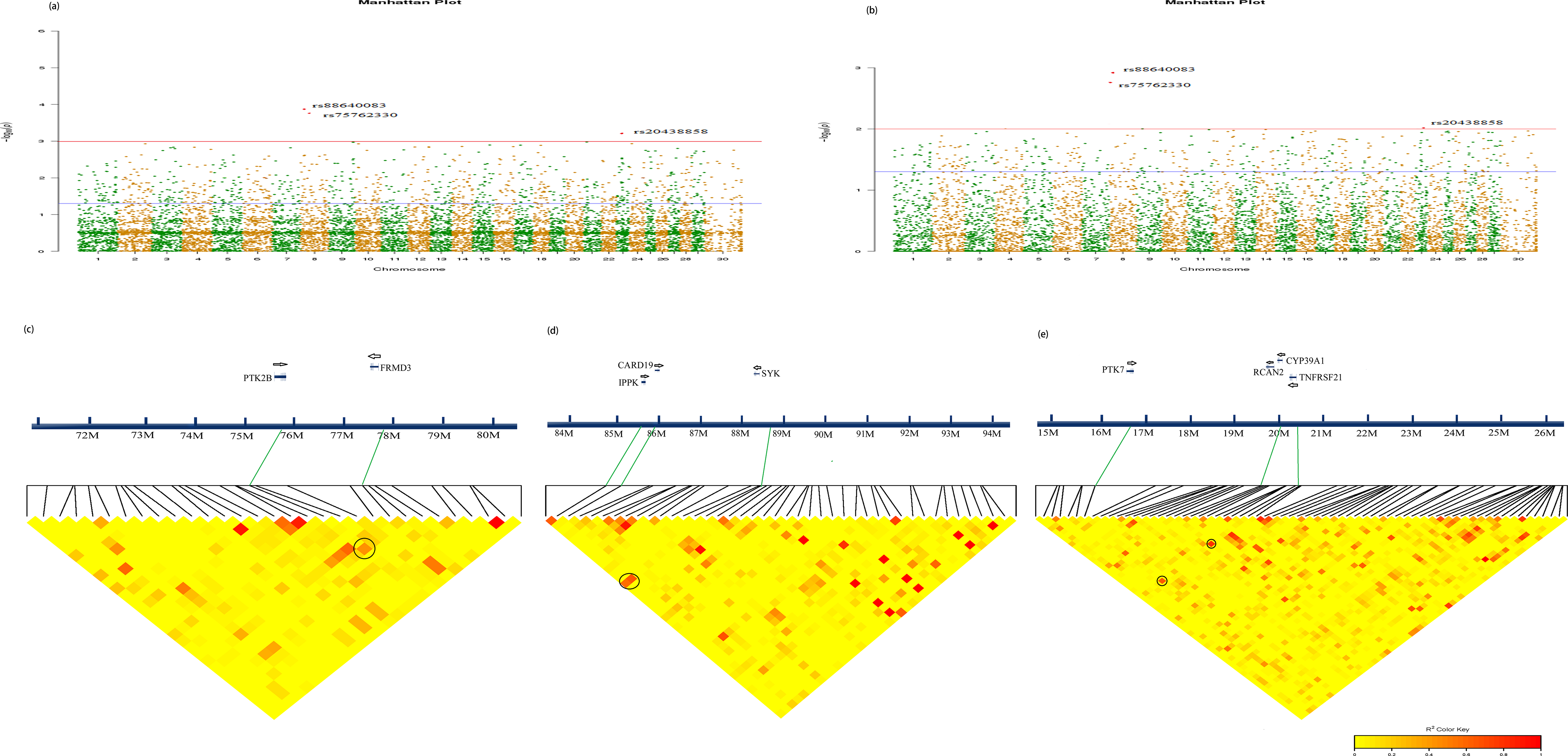
Three Significant SNPs associated with genes chromosome mapping for Chinese Holstein. Manhattan plots (a) and (b) showed the SNPs associated with mastitis in Chinese Holstein screened by two models, respectively. (a) was the result of Bayesian analysis, while (b) represented the related genes labeled by logistic analysis model. Red dots represented the chromosome location of the associated genes. (c-e) were partial LD block of three significant SNPs, respectively, with a distance interval of 1Mb, the more reddish the LD block color is, the stronger the correlation the dots. SNPs rs75762330 and rs77816736; rs88640083, rs85927029 and 8563916; and rs20438858, 19736020 and rs16711445 were in the same LD block (black circle), respectively, suggesting that their corresponding genes potential association with each other, respectively.

### Correlation between SNPs

Calculated LD coefficients between two pairs of SNP markers in the genome, and then LD coefficients classified according to the distance between the markers (Figure 4). Finally, the average LD coefficients between molecular markers at a certain distance counted. The average LD coefficient of 100kb on the genome of case-control two Chinese Holstein cows was about 0.5. However, the corresponding LD coefficient was still above 0.3 when the distance was 1000 kb. Moreover, we could also observe from the figure that the LD decay speed and C value were same between case-control groups. Of course, we also noticed that LD decayed very slowly. 2b-RAD data shows that our SNP markers are sparse (9589 bp). Therefore, a suitable LD block map gained when the classification interval size was set to 10Mb.

**Figure 4.**
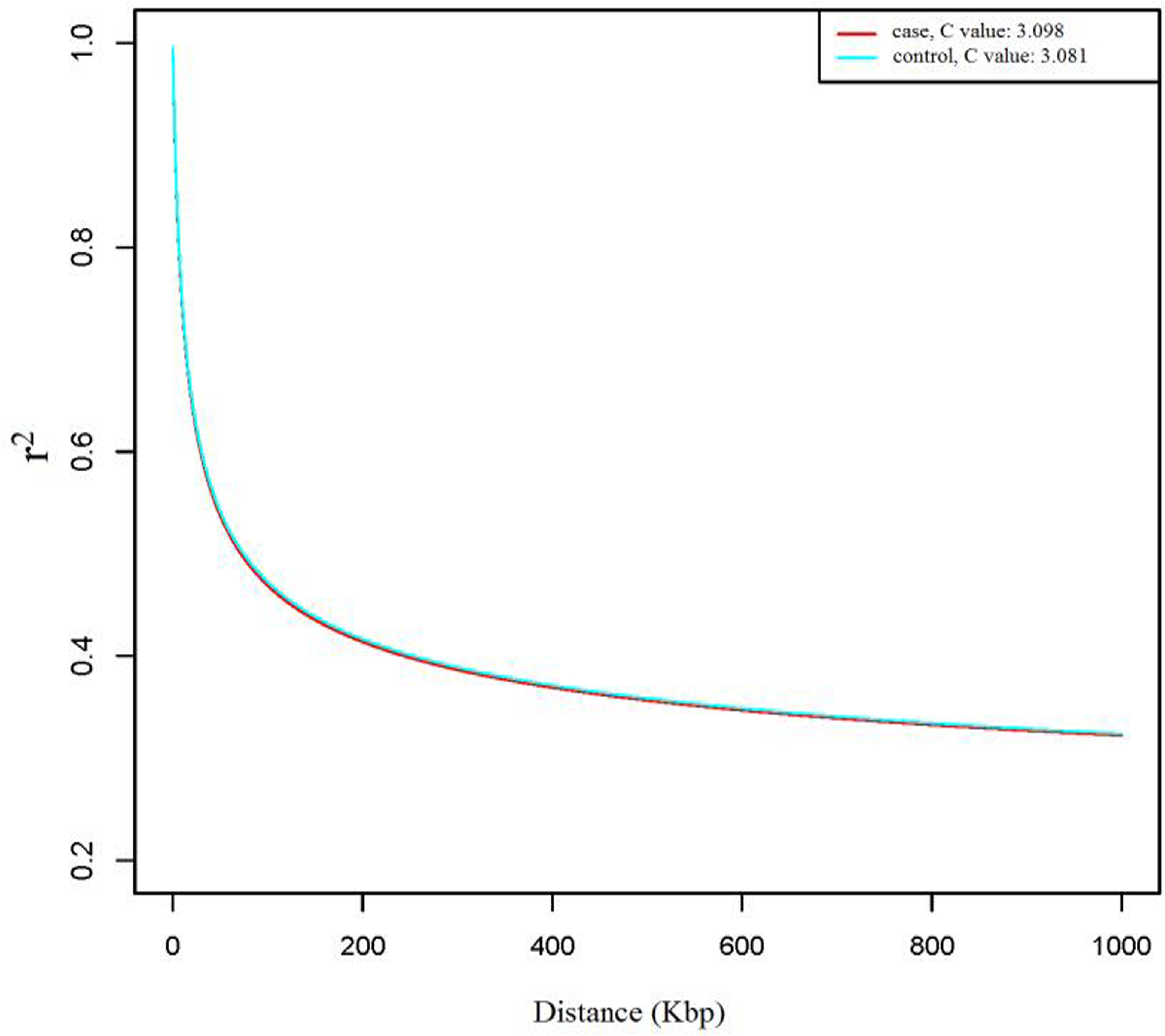
The Linkage Disequilibrium attenuation curve of case-control SNPs. The lines of different colors represented different populations/chromosomes, horizontal coordinates were the physical distances between the SNPs pairs, and vertical coordinates were the average r2 values of the same physical distance marker pairs. As the distance between sites increases, r2 usually showed a decreasing trend. The larger the C value, the lower the probability of recombination between SNPs and LD attenuation distance; the smaller the C value, the higher the probability of recombination between SNPs and LD attenuation distance.

Figure 3(c) showed that rs75762330 associated with rs77816736, however, the latter P-value was >0.05 in both analytical models (Table S6). SNPs rs88640083 associated with rs85927029 and rs85635916 (Figure 3(d)), yet the latter two were statistically meaningless. SNPs rs19736020 and rs16711445 associated with rs20438858 (Figure 3(e)), and Table S6 shown that the first two were not statistically significant. We also calculated the linkage disequilibrium (LD) between three significant SNPs. Genetic linkage analysis showed that SNPs rs75762330 not correlated with rs88640083 (r^2^= 0.0022) and rs20438858 (r^2^= 0.043). However, rs20438858 weakly correlated with rs88640083 (r^2^= 0.22).

### Three significant SNPs population verification

Correlation analysis performed on three important SNPs in another larger independent Chinese Holstein dairy population via direct sequencing (Figure 5, Table 4). We successfully performed PCR cloning near three important SNPs, then direct sequencing. Data shown that the three locus’ P-value was <0.05, indicated that they were statistically significant associations between Chinese Holstein cow mastitis. The correlation between rs20438858 and risk of mastitis was still statistically significant, with the adjustment allele OR= 0.359（OR <1）. While, the other two significant SNPs (rs75762330 and rs88640083) located on BTA 8, with the adjustment allele OR = 2.416 and 1.879 (OR >1), respectively. Table 4 also shown that base G in rs88640083 had higher occupancy rate in case group than control group. And in rs75762330 base T as well. However, in rs20438858, the probability of base T in case group was less than control group. We also noted that AF*e* value for rs75762330 and rs88640083 was 0.2489 and 0.2426 >0, respectively; while rs20438858 AF*e* value was −1.786<0. Three significant SNPs annotated to three candidate genes

**Figure 5.**
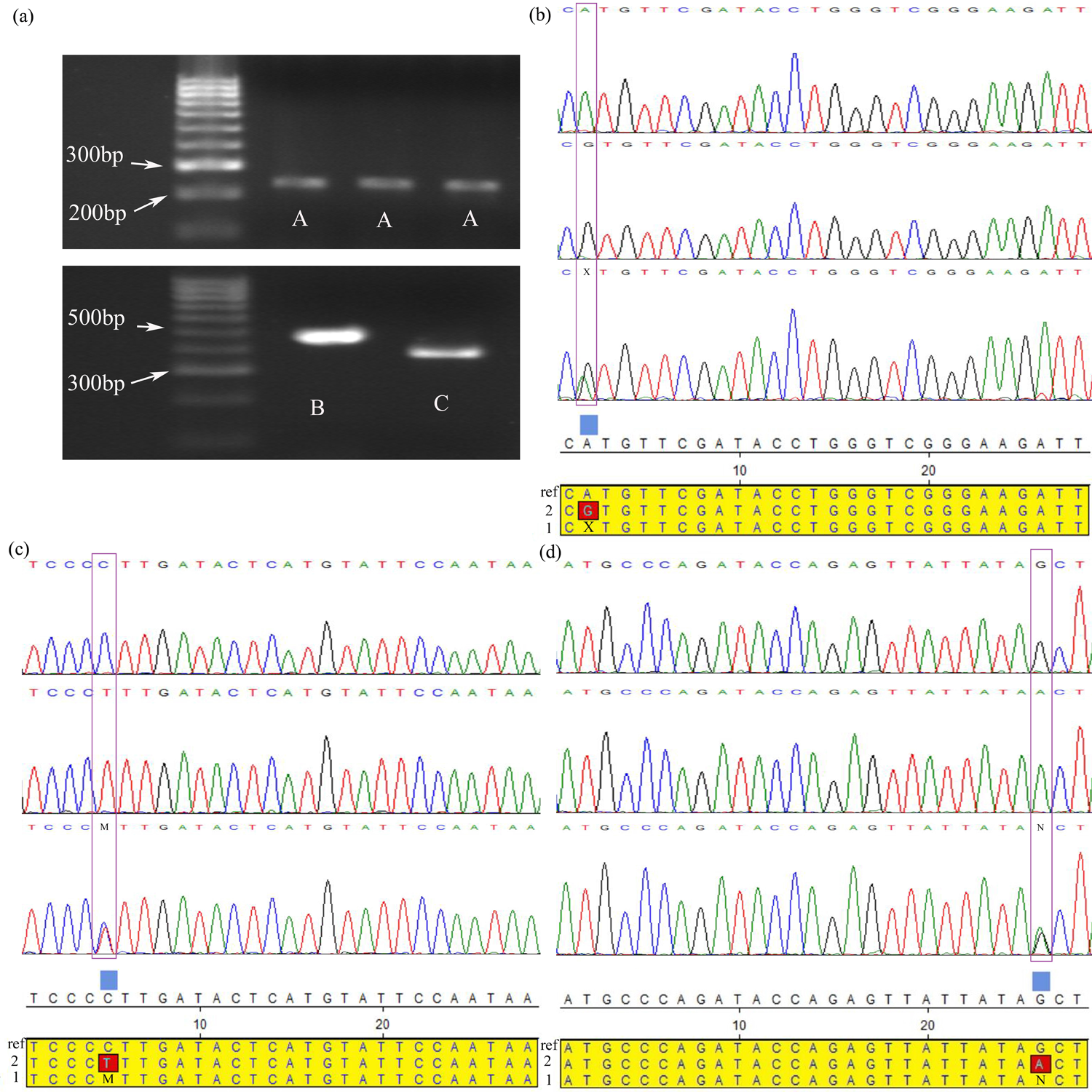
Three SNPs were directly sequenced in an independent validation population of Chinese Holstein cows. (a) Gel electrophoresis pattern PCR amplified fragments near three significant SNPs, A-C were PCR amplified fragments of SNPs rs88640083, rs75762330 and rs20438858 regions, respectively. (b-d) Directing sequencing results of PCR amplification products near above three important SNPs, and their alignment with reference sequences (ref: reference sequences; 1: heterozygous sequences; 2: variant sequences). The purple boxes were where the three SNPs located. X, M and N represented the heterozygous types of the three SNPs, respectively.

GO enrichment analysis indicated that three important genes associated with adaptive and innate immune response in Chinese Holstein cows (Figure S6). Figure 6 also showed that these three candidate genes directly or indirectly affected the function of AKT1 and promoted the expression of pro-inflammatory cytokines in mammary epithelial cells and macrophages of dairy cows, suggested that these three genes are involved in mammary epithelial cells and macrophages the polarization-related biological functional activities. AKT1 (protein kinase B), As a key Jak2/STAT5 pathway protein, plays an important role in the regulation of differentiation, secretion, survival and proliferation of mammary epithelial cells and also plays a key role in mammary remodeling and lactation sustainability in dairy cows (MAROULAKOU *et al.* 2008; CHEN *et al.* 2010; CREAMER *et al.* 2010; ARRANZ *et al.* 2012; HOU *et al.* 2016), which is bound to play a key role in mediating the immune response to mastitis in dairy cattle.

**Figure 6.**
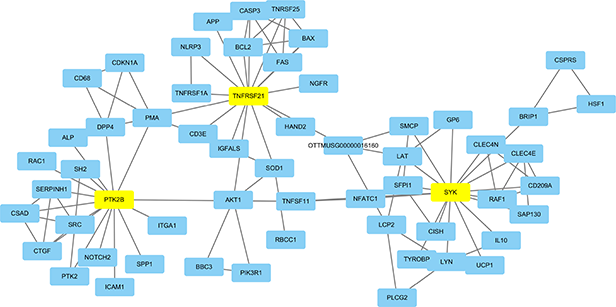
Candidate genes interaction network Diagram based on KEGG Database. The network map was constructed with the three candidate genes as the core. The interaction between genes was represented by a line.

## Discussion

Genetic analysis of GWAS had a considerable impact on the study of complex genetics (LOH *et al.* 2015). GWAS has also achieved unprecedented success in identifying gene regions and candidate gene variants closely related to clinical phenotypes and disease susceptibility (chromosome and gene level, in term of the association between SNPs and traits) (YANG *et al.* 2011; LEE *et al.* 2012; CROSS-DISORDER GROUP OF THE PSYCHIATRIC GENOMICS *et al.* 2013). GWAS develops new functional studies and provides therapeutic strategies by comparing multiple gene regions or candidate genes, identifying new candidate genes for causal pathways (DEELEN *et al.* 2013). Moreover, identifying associated gene mutations could help reveal the pathogenesis of disease and provided cut-in points for treatment, and analysis of common genetic variations identified many risk loci for multiple complex diseases (XU *et al.* 2012; BERNDT *et al.* 2013). However, knowledge of disease biology and treatment remains limited. Gene’s functional changes caused by mutations associated with cow mastitis, which are also subtle and difficult to explain. Two-stage correlation analysis for three mastitis significant SNPs

Genomic prediction methods of genetic values might show different results for different phenotypes, and the results might be different due to different genetic structures among traits (MOSER *et al.* 2009; RESENDE *et al.* 2012). To improve accuracy, we used two-stage association analysis to reduce false positives. We reduced the dimensions when processing data, considering only the SNPs associated with inflammation and immune response. In stage I, considering that case-control state in the present study was not in a normal distribution, and to obtain accurate mastitis significant SNPs and genes information in Chinese Holstein, we tried to use two analysis models (Bayesian model and logical regression model) to carry out GWAS of 2b-RAD sequencing results. Several genetic background analyses have been found related to mastitis traits in dairy cattle populations, although to our knowledge no study has been conducted in Chinese Holstein dairy population to date using two GWAS analysis models at the same time. Comparison between the two models showed that although there were differences in SNP tagged under the same P value (P<0.05), the general trend of the association with mastitis was similar. The results suggested that Bayesian screen out more accurate significant SNPs (42, P<0.001, Table S4) in dairy cows, while logical regression analysis identified more SNPs (51, P<0.01, Table S5). Importantly, we identified three (rs75762330, rs88640083 and rs20438858) novel dairy cows mastitis traits significant SNPs in Chinese Holstein cows. SNPs rs75762330 within PTK2B and SNPs rs88640083 in intergenic nearby SYK located on BTA 8 were risk factors (OR>1), and the SNPs rs20438858 (OR<1) in TNFRSF21 located on BTA 23 was a protective factor for dairy cows mastitis.

With regarding to stage II, we used a case-control study to verify the association of three important SNP markers with cow mastitis. It compared the exposure ratios of important SNPs in case and control groups (BAGHERI *et al.* 2016; WEISSBROD *et al.* 2018; ZHOU *et al.* 2018). After statistical test, if there is significant difference between two groups, it can be considered that the SNPs associated to cow mastitis. When comparing the two groups, excluded the interference from external matching factors and only considered the relationship between SNPs and mastitis. According to the Pitman efficiency increment formula (2R/(R+1)), determined the appropriate sample size and gained higher test efficiency. Here, we validated the association of three important SNPs with cows mastitis. SNPs rs75762330 and rs88640083 were correcting factors (AF*e*>0) for cows’ mastitis, which associated with mastitis susceptibility. SNPs rs20438858 as a negative regulator (AF*e*<0) for cows’ mastitis and associated with mastitis resistance.

### Three significant SNPs are located in genomic non-coding sequences

Previous studies found that conserved non-coding regions (CNCs) in introns and near genes show large allelic frequency shifts, similar in magnitude to missense variations, suggesting that CNCs are critical for gene function regulation and evolution in many species, including yeast, fruit flies and vertebrates (HAUDRY *et al.* 2013; VISSER *et al.* 2014; PETIBON *et al.* 2016; DICKEL *et al.* 2018). However, The CNCs variation, which does not directly change the amino acid sequence, is the key to the regulation of gene genetic information expression and affects biological functions and diseases in mammalian (PATRUSHEV and KOVALENKO 2014). Our GWAS data provided a statistical list of SNPs associated with mastitis traits in dairy cows, where the associated significant SNPs are located in non-coding regions (intron and intergenic) of the genome. Functional annotations showed that the three SNPs (rs75762330, rs88640083 and rs20438858) were associated with immune and inflammatory responses in dairy cows, implicating them as key SNPs for mastitis in dairy cows. But the biological function behind this statistical association is still not known, because this association may stem from hindering another biological function, such as regulating function, or being affected by other functional SNPs, and this can only be illustrated by subsequent experimental studies.

### Significant SNPs are at low or moderate genetic polymorphisms

Pathogen-specific mastitis traits are a direct indicator of cow mastitis infection. GWAS studies showed that the mastitis trait is a low genetic polygenic trait that is controlled by multiple sites distributed in the genome, and the genetic effect of each locus is relatively small (WU *et al.* 2015). Our data results were basically consistent with previous studies. Stage I data showed that rs88640083 (PIC=0.1676<0.25) was low polymorphism, while rs75762330 (0.25<PIC=0.2999<0.5) and rs20438858 (0.25<PIC=0.3366<0.5) were moderately. Three SNPs associated with mastitis also demonstrates that cow mastitis has multiple genetic effects.

### Three important candidate genes biological function

Innate immune system is a key protective mechanism of bovine mammary gland against exogenous pathogen infection. GO function analysis annotated three significant SNPs into three important genes (PTK2B, SYK and TNFRSF21), which suggested that these three genes are novel candidate genes associated with mastitis traits in Chinese Holstein cows. PTK2B involved in regulating the LPS-TLR4 cascade in macrophages and affected the migration of dendritic cells (DCs) (RACIOPPI *et al.* 2012; RHEE *et al.* 2014). It was also an important homeostasis regulator in natural immune cells such as bone marrow mononuclear cells (RACIOPPI *et al.* 2012; RHEE *et al. 2014*; LLEWELLYN *et al.* 2017). As for SYK, it played an essential role in signal transduction of adaptive immune receptors and participated in the regulation of innate immune recognition, vascular development, platelet activation and cell adhesion (MOCSAI *et al.* 2010). Studies reported that the SYK was also involved in regulating the proliferation of dairy mammary epithelial cells, affected milking cycles and milk production (HOU *et al.* 2016). TNFRSF21 might play an important role in regulating the degeneration of the mammary gland and providing protection against infection (KHALIL *et al.* 2011). We also noted that SYK and TNFRSF21 involved in “Toll-like” and “TNF/TNFR” signaling pathways, respectively, which are the key pathways to identify exogenous pathogens and induce inflammation and immune response.

## Conclusions

In this study, we committed to improve understanding biogenetic variation of mastitis in Chinese Holstein cows, and to guide the construction of ant-mastitis populations and improve the populations’ anti-mastitis characteristic in dairy cows. Therefore, reduced-representation sequencing (2b-RAD) used to systematical study the conventional genetic variation (direct genotyping) of Chinese Holstein cows. And then rely on two-stage correlation analysis to find significant SNPs associated with risk of mastitis. Finally, we screened out three significant SNPs (rs75762330, rs88640083 and rs20438858) associated with immune response and inflammation, which suggested that these three genes (PTK2B, SYK and TNFRSF21) are novel candidate genes associated with mastitis traits in Chinese Holstein cows.

### Abbreviations

2b-RAD, type IIB endonucleases restriction-site associated DNA; GWAS, Genomic wide association studies; SNPs, single nucleotide polymorphisms; PTK2B, protein tyrosine kinase 2; SYK, spleen tyrosine kinase; TNFRSF21, tumor necrosis factor superfamily member 21; TLR4, Toll-like receptor 4; NF-κB, nuclear factor-kappa B; PCA, principal component analysis; PC, principal component; Q-Q plots, Quantile-Quantile plots; OR, Estimated odds ratio; PIC, Polymorphism Information Content; He, Heterozygosity Expectation; Ho, Heterozygosity Observation.

### Ethics approval and consent to participate

This experimental animal and its care protocol followed previous studies. National and local animal welfare agencies (Jiangsu province, China; Nanjing Agricultural University and Nanjing Weigang Dairy Co., Ltd; Approval No.20160615) approved all experimental animal procedures in this study.

### Consent for publication

The authors announced their agreement to publish the manuscript

### Availability of data and materials

The entire genome reference sequence raw data is based on the sequence provided by the NCBI database (ftp://ftp.ncbi.nlm.nih.gov/genomes/all/GCF/000/003/055/GCF_000003055.6_Bos_taurus_UMD_3.1.1/GCF_000003055.6_Bos_taurus_UMD_3.1.1_genomic.fna.gz).

Summary information and download links for all SNPs (ftp://ftp.ncbi.nlm.nih.gov/genomes/all/GCF/000/003/055/GCF_000003055.6_Bos_taurus_UMD_3.1.1/GCF_000003055.6_Bos_taurus_UMD_3.1.1_genomic.gff.gz).

Supplementary data are supported by diagrams and tables as powerful data for manuscripts.

## Authors’ contributions

YC, JL, HL and GW co-designed the experiment. FC, LL, CL, DY, JC, CX and JL participated in the screening of experimental animals and blood sample extraction. FY and FC sorted and analyzed 2b-RAD sequencing data. FY and TB wrote and calibrated experiments manuscripts. The final manuscript was read and authorized by all authors.

## Acknowledgements

We thank Nanjing Weigang Dairy Co., Ltd. for providing experimental Chinese Holstein blood samples and Shanghai Oe Biotech Co., Ltd. for providing 2b-RAD genome sequencing technology support.

---

Table 1 Significant SNPs screened by the Bayesian Model and Logical regression analysis model

Note: * indicated the P-value calculated by Chi-square (<0.001); ** is the t-statistic P-value of the logical regression model (<0.01);

CHISQ is Chi-square under Chi-square test. STAT is the t-statistic coefficient under the logistic regression model.

OR: Estimated odds ratio. L95: Lower bound of 95% confidence interval for odds ratio. U95: Upper bound of 95% confidence interval for odds ratio. Nan: meaningless number. Na: missing value or Not available.

Table 2 Significant SNPs genetic diversity and their Go enrichment annotations

Note: He: Heterozygosity Expectation. Ho: Heterozygosity Observation. PIC: Polymorphism Information Content.

Table 3 GO enrichment items and genes of three significant SNPs.

Table 4 Case-control study analyzed three significant SNPs in independent validation population.

## Additional files

### Additional file 1

Figure S1. A 2b-RAD sequencing Diagram for Chinese Hesitant cows.

Figure S2. Base distribution map (a) and base mass distribution map (b). (a): the base position of Reads is represented by the horizontal coordinate, and the ordinate is the proportion of the base; different colors represent different base types, and the unrecognized base in sequencing was N; the base distribution of reads at the R1 end were located at the left of the dotted line at 150bp; the right 150bp was the base distribution of R2 terminal reads. b: the lateral coordinates represented the base position of reads, and the ordinate was the mass value of the base at the corresponding position; the base mass value of Double terminal sequencing reads at the R1 end were located at the left of the dotted line at 150bp; the right 150bp were the distribution of base mass value of reads at R2 terminal; the darker the blue color, the higher the base ratio of the mass value in the data.

Figure S3. Label distance distribution histogram. The horizontal axis represented the distance between the two adjacent labels; the vertical axis represented the number of the labels corresponding to the distance; the upper right was the box line diagram of the adjacent labels of the labels, and the horizontal line in the middle of the box line chart was the median, that is, the average spacing between the labels.

Figure S4. The Distribution of SNPs on Chromosomes in Chinese Holstein dairy cows. The horizontal axis indicated the coordinates of the physical position of the chromosomes; the vertical axis represented the number of the corresponding SNPs (window size, 20Kbp; Step length, 10Kbp).

Figure S5 Principal Component Analysis and RAD typing of SNPs. (a) Abscissa represented principal component 1 (PC1); ordinate represented principal component 2 (PC2); each point was a sample with different shapes and colors representing different groups. (b) Genotyping and genotype score of 10058 SNPs. Abscissa: 0 (two bases of the type were different from the reference genome); 1 (two bases of typing were the same as one of the reference genomes); 2 (two bases of typing were the same as those of the reference genome).

Figure S6 Hierarchical Network of candidate Gene function based on go enrichment Analysis. Each circle represented a Go entry; the color indicated the enrichment degree, the deeper the color (yellow), the more genes enriched in the Go entry; the direction of the arrow indicated hierarchic relationship.

### Additional file 2

Table S1. Illumina base recognition and mass value correspondence table

Table S2. The top 5 principal component data of all SNPs for each sample

Table S3. Three genotypic risk estimation data of case-control by Armitage-test

Table S4. Forty two important SNPs screened out via Bayesian Analysis Model.

Table S5. The logistic regression model screened out 51 important SNPs.

Table S6. Five SNPs screened by the Bayesian Model and Logical regression analysis model.

